# Categorising update mechanisms for graph-structured metapopulations

**DOI:** 10.1101/2022.10.20.513032

**Authors:** Sedigheh Yagoobi, Nikhil Sharma, Arne Traulsen

## Abstract

The structure of a population strongly influences its evolutionary dynamics. In various settings ranging from biology to social systems, individuals tend to interact more often with those present in their proximity and rarely with those far away. A common approach to model the structure of a population is Evolutionary Graph Theory. In this framework, each graph node is occupied by a reproducing individual. The links connect these individuals to their neighbours. The offspring can be placed on neighbouring nodes, replacing the neighbours – or the progeny of its neighbours can replace a node during the course of ongoing evolutionary dynamics. Extending this theory by replacing single individuals with subpopulations at nodes yields a graph-structured metapopulation. The dynamics between the different local subpopulations is set by an update mechanism. There are many such update mechanisms. Here, we classify update mechanisms for structured metapopulations, which allows to find commonalities between past work and illustrate directions for further research and current gaps of investigation.

## 1 Introduction

The spatial structure of a population has a considerable impact on the evolutionary dynamics of a population. One of the most popular theories for studying the effect of underlying structure on the evolution of a population is Evolutionary Graph Theory [1], where graphs represent spatial structures. The nodes of a graph represent individuals, and links indicate individual’s neighbours. A link determines where an individual can place their offspring. An update mechanism describes which individuals produce offspring and how this offspring is placed. Here, we focus on fixed structures, but the framework can be extended to the case where a spatial structure itself varies over time [2–5].

An extension of evolutionary graph theory is given by graph-structured metapopulations in which the nodes indicate subpopulations, and the links indicate the migration between subpopulations. Given that fragmented habitats are ubiquitous in ecology, metapopulations, and their evolution and ecology have been extensively studied [6–8]. However, there are few studies on graph-structured metapopulations with more complex graphs [9–11] and thus, this issue calls for further investigation. Depending on the system, different update mechanisms have been applied to describe the dynamics. In general, evolutionary dynamics are not robust to the choice of update mechanisms [5, 12, 13]. An important factor that changes the system’s dynamics is how selection acts. Depending on the update mechanism, selection can be global or local. This will affect the evolutionary dynamics dramatically. The two most important events that govern the dynamics of the population are birth and death. The order in which birth and death occur, which often determines if selection is local or global, has a high impact on the fate of the population [14].

We aim to categorize the different update mechanisms for the evolution of structured metapopulations to facilitate future work. First, we recall the update mechanisms on graphs of individuals, and then we generalize them to graphs of subpopulations.

## 2 Update mechanisms in graphs of individuals

In graphs of individuals, three main events influence the evolution of a population: birth, mutation, and death. In the long run, a mutation-selection balance is reached [15–18]. Here, we focus on the low mutation regime in which the system reaches fixation or extinction of the mutant before the next mutation occurs. In that case, the population is typically homogeneous, and mutations reach fixation one after another. In the low mutation regime, the two quantities of most interest are the fixation probability and the average time to fixation.

If the fixation probability for advantageous mutants on a graph is higher than the fixation probability of the complete graph (and vice versa for disadvantageous mutations), the graph is called an amplifier of selection. On the other hand, if the fixation probability for advantageous mutants on a graph is less than the fixation probability on the complete graph (and vice versa for disadvantageous mutations), this graph is called a suppressor of selection. These notions have been first introduced in [1].

The order of birth and death events and how selection acts upon them can substantially influence the population’s fate. We use the following scheme to differentiate between the update mechanisms. For birth, we use **b** if birth is random, i.e., a birth-giving individual is chosen randomly, and **B** if the birth-giving individual is chosen with probability proportional to its selection parameter for birth. Similarly, we represent the death event by **D** if the individual is selected for death with a probability associated with its selection parameter for death and by **d** if the individual dies at random.

Accordingly, the possible update mechanisms are **BD**, **Bd**, **bD**, **bd**, **DB**, **Db**, **dB**, and **db**, where the order shows which event is first, see Table 2.1. In all these update mechanisms, the first event is global, meaning that the individual is selected from the whole population. In contrast, the second event is local because the individual is selected from a subset of the population in the neighbourhood of the first individual.

**Table 2.1:**
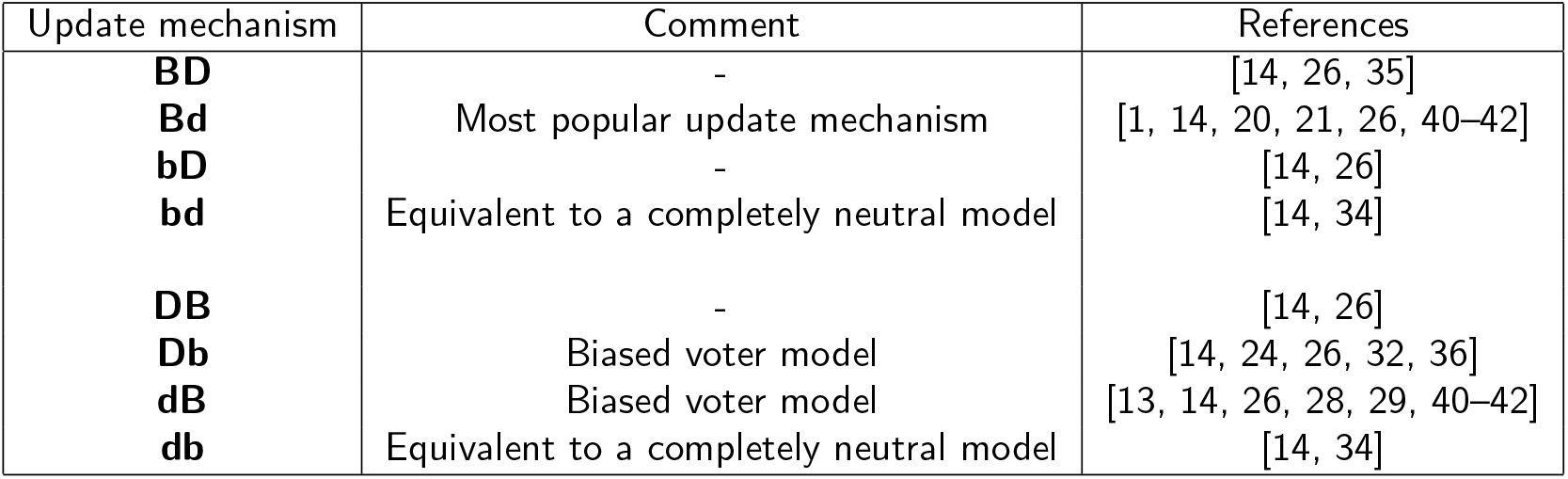
Update mechanisms in graphs of individuals. In these update mechanisms, birth and death change the state of the population. The first event is global and the second event is local.

As an example, applying the update mechanism **Bd** in a well-mixed population of size *N* consisting of two types of individuals, *N* – *n* wild-type and *n* mutants, with mutants having a relative selection parameter for birth *r* with respect to wild-types. To increase the number of mutants in the population, one mutant is selected for reproduction (global event). Then, from the neighbours of the mutant, one wild-type is selected for death (local event). The offspring fills the empty spot of the dead individual. The probability of increasing the number of mutants by one is then

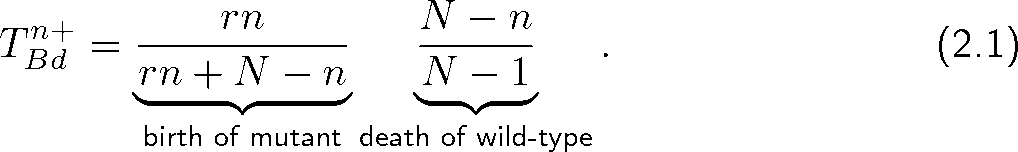

Similarly, the probability of decreasing the number of mutants by one is

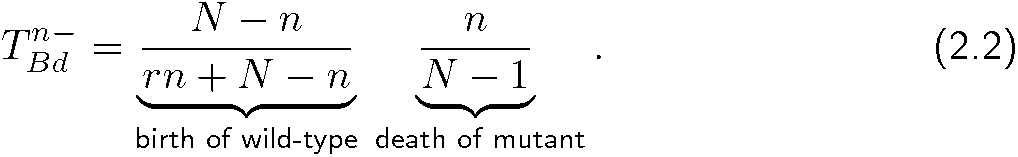

Using the recursive relation for the fixation probability [19, 20], the fixation probability starting from *n* mutants is

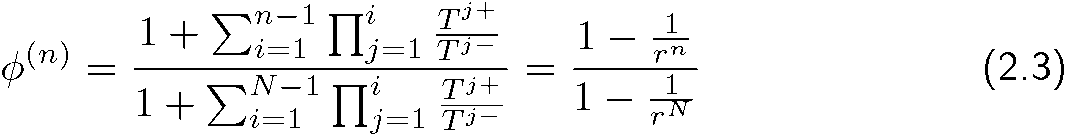

The update rule **Bd** has been vastly explored in both structured and well-mixed populations [1, 21–26]. For small populations under the update mechanism **Bd**, most small structures are amplifiers of selection, and only a minimal fraction of structures suppress selection [5]. In larger populations, in the weak selection regime, there are quite a lot of graphs that suppress selection [27]. Furthermore, some structures have the same fixation probability as the complete graph. This is the case when the total weight of incoming edges to a node is homogeneous through the entire graph, as shown in the “isothermal theorem” [1, 20].

Similarly, many studies are dedicated to **dB** updating [5, 13, 14, 22, 23, 26, 28–31]. Contrary to **Bd**, only a small fraction of undirected graphs under the update mechanism **dB** amplify selection and the majority suppress selection [5]. Another popular update mechanism is **Db** which is the same as the score-dependent viability model introduced in [32], this is equivalent to the voter model in statistical physics [24, 25, 33]. In [26], it is shown that the dynamics of a lattice of an integer dimension under update mechanisms **Bd** and **Db** are equivalent. It is illustrated however, that the dynamics of this lattice under update mechanisms **dB** and **bD** are fundamentally different.

Among the 8 update mechanisms for graphs of individuals, **bd** and **db** are identical, describing a system where natural selection has no role in the evolution of the population [14, 34]. In the update mechanisms **DB** and **BD** selection acts both on death and birth [14, 26, 35], but they are not equivalent in general. Intuitively, it is expected that the fixation probability of an advantageous mutant is higher in the presence of update mechanisms **DB** and **BD** than in the other update mechanisms. The general transition probabilities for these 8 update mechanisms are given in Appendix A. In the Fig. 1, the transition probability for a specific example under the 8 update mechanisms is given.

**Figure 1:**
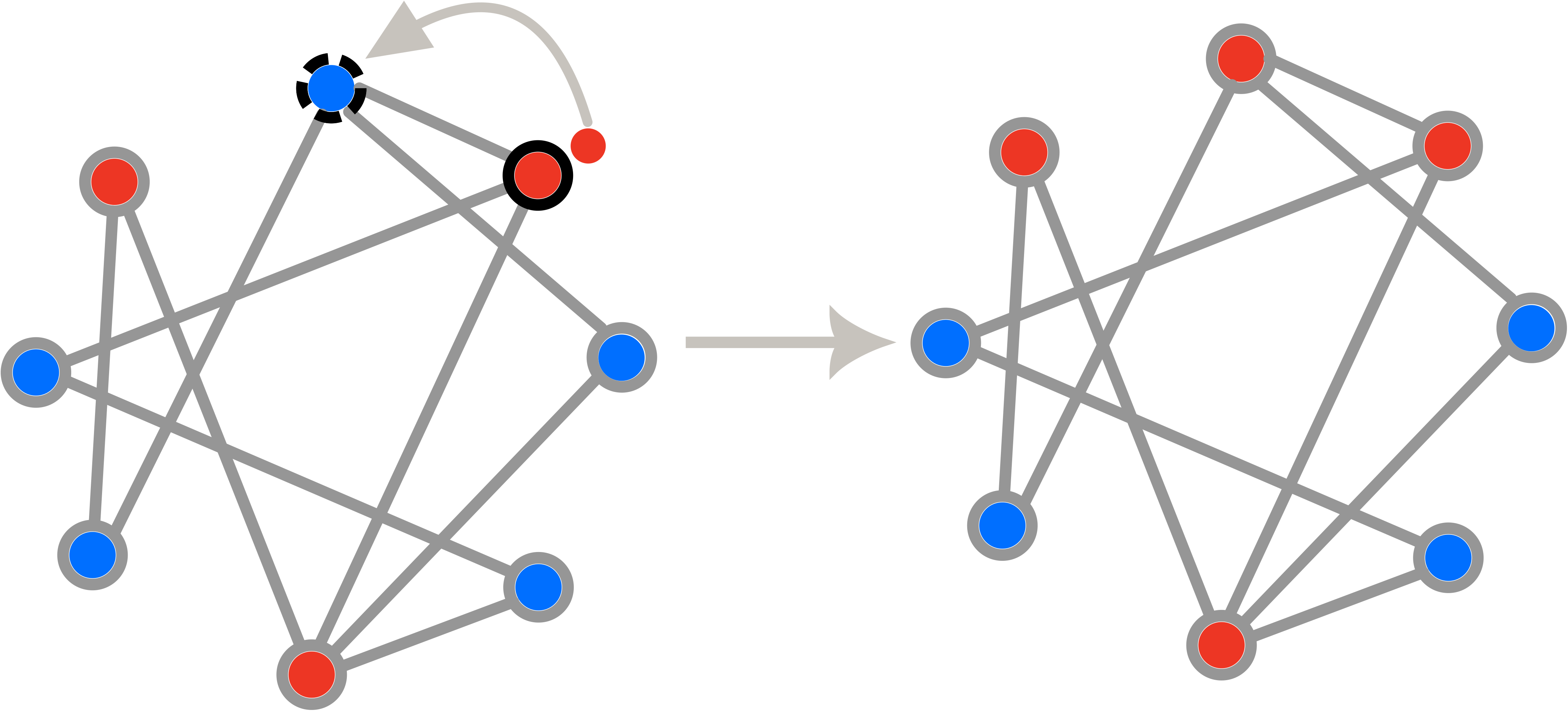
Different update schemes for graph of individuals. We consider a random population structure of size eight with five wild-type individuals (in blue) and three mutant individuals (in red). Neighbours are connected via links. Individual encircled with solid black circle represents the birth giving parent, whereas, the individual encircled with black dashed circle is the one chosen for death. The population size remains constant throughout the dynamics with offspring replacing dead individuals. Assuming that the selection parameter for birth of a mutant individual is *r* = 2 (1 for the wild type) and that the selection parameter for death of the mutant is *t* = 1/2 (1 for the wild type), the probabilities that the transition shown in the figure takes place, is different for the different update mechanisms shown in Tab. 2.1. For example, in the case of **BD**, the probability to choose this particular mutant individual for birth is 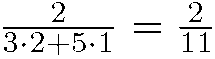. The probability to choose this particular wild type neighbor for death is 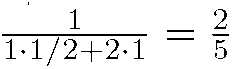, which leads to a probability 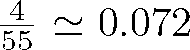. Similarly, we find: **Bd**: 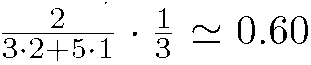. **bD**: 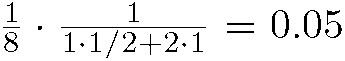. **bd**: 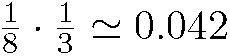 **DB**: 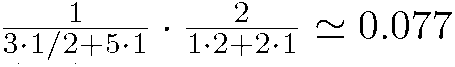. **Db**: 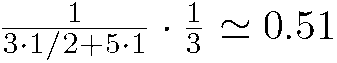. **dB**: 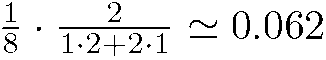. **db**: 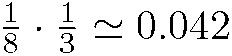.

In addition to the above update mechanisms, there are other update mechanisms in which, instead of two individuals (one for birth, one for death), one edge is selected [24, 36, 37]. Then an individual dies at one end, the other gives birth, and the offspring fills the neighbouring empty spot. In this update mechanism, selection can act on death and birth events or not. Here, we will not consider this kind of update mechanism.

For more information about the comparison of different update mechanisms on various graphs with different features, we refer to the following references: In [36] the authors ask when the fixation probability in an evolutionary graph equals the fixation probability in a Moran process. A Moran process [38] is equivalent to the update mechanism **Bd** in a complete graph. In [39], the authors investigate the evolutionary game dynamics on the star graph in the presence of different update mechanisms. They show that the evolutionary dynamics of heterogeneous graphs is not robust under the choice of update mechanism. In [37] the effect of the directionality of a graph on its evolutionary dynamics for different update mechanisms is investigated. It has been shown that regardless of the update mechanism, the directionality always suppresses selection. In this manuscript, we extend the above update mechanisms to graphs of subpopulations where each node comprises a well-mixed subpopulation. The links indicate the migration between the patches (see Fig. 2).

**Figure 2:**
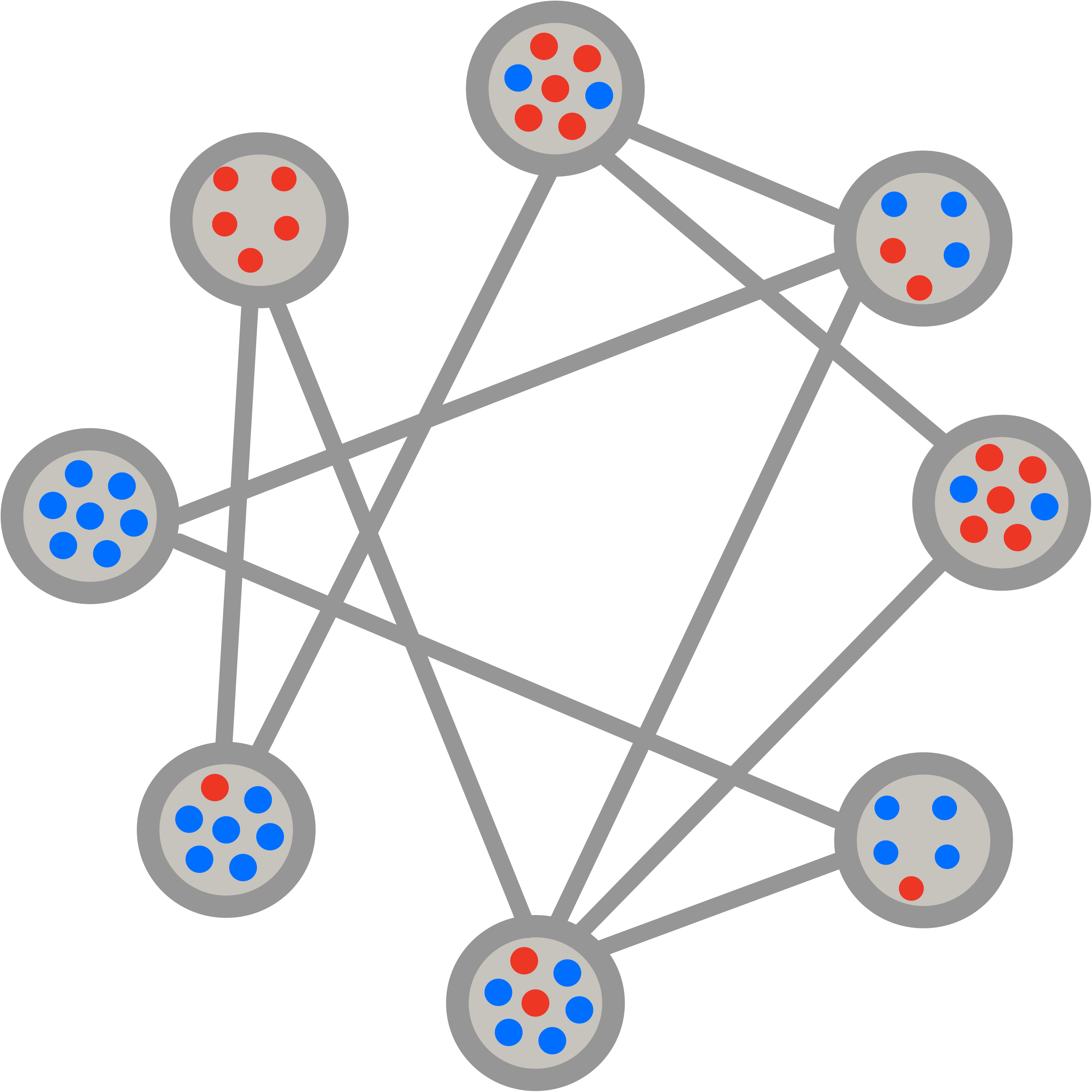
Graph of subpopulations. Each node in the graph includes a well-mixed subpopulation and each link indicates migration between two subpopulations.

## 3 Categorising update mechanisms for graphs of subpopulations

In most of the potential applications of evolutionary graph theory, including biological and social systems, individuals tend to interact locally in subpopulations, with some rare interactions with other groups of individuals. This is because the populations are segregated for various reasons, and they are geographically distant. Individuals in the same geographical area compete over resources or provide common goods. However, there is an occasional migration to and from other geographical areas. In [10], it has been shown that for a particular update mechanism, the results of evolutionary graph theory [1] do not carry over into graphs of subpopulations. For example, a star-structured metapopulation does not always amplify selection — in some migration regimes, it suppresses selection. It will be interesting to see if this is the case for other update mechanisms.

In a model that retains the local population’s size fixed, birth and death either happen in the same subpopulation, or the first event can be followed by an individual’s migration to or from another subpopulation. Having this in mind, we can categorize update mechanisms into two groups:

- A set of update rules where there is always the possibility that the first event is accompanied by migration (Fig. 3 A-B).
- A set of updates where birth and death events are always in the same subpopulation and migration happens independently from birth and death, (see Fig. 3A & C).

**Figure 3:**
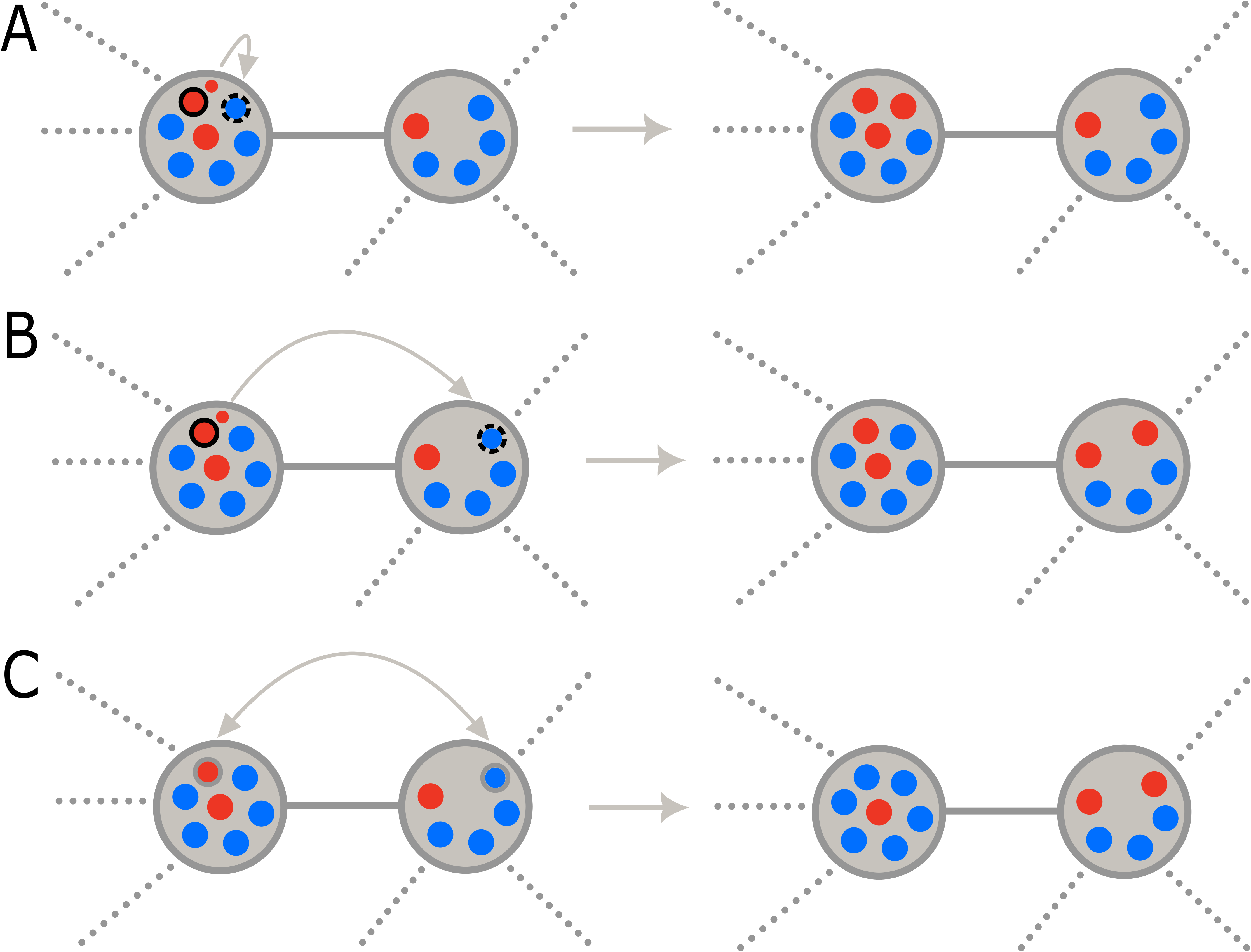
Update mechanisms in a graph-structured metapopulation. The population consists of two types of individuals, wild-types (blue) and mutants (red). The individual marked with the solid black line gives birth and the individual marked with black dashed line is selected for death. (A) *Birth-death or death-birth in one patch without migration:* This includes birth-death or death-birth in a coupled update mechanism without migration as well as an uncoupled update mechanism in which both death and birth happen in the same subpopulation. (B) *A coupled update mechanism with migration:* if birth is coupled with migration after each birth the newborn migrates to an adjacent patch and replace one of its individual. If death is coupled to migration a death in one patch is followed by birth of an individual in one of the adjacent patch where newborn occupies the place of dead individual. (C) *Migration in an uncoupled update mechanism:* migration happens independently from birth or death and it only exchanges the position of two individuals from different patches. This process is completely random.

We refer to the former category as update mechanisms with coupled migration and the latter as update mechanisms with uncoupled migration. The order of events (birth and death) in each of these classes and how selection acts upon them might affect the dynamics considerably. In both categories of update mechanisms, selection for the first event can act on both patch and individual levels, i.e., one first selects a patch and then an individual from the chosen patch. Selection on the second event can act both on the patch and individual levels in the update mechanisms with coupled migration. However, in the update mechanisms with uncoupled migration, selection in the second event always acts on the individual level since the second event happens in the same patch as the first event.

We code the update mechanisms as follows: the first letter stands for migration to show if it is coupled (**M**) or uncoupled (**m**). If selection on an event acts both at the patch and individual levels, we allocate two labels to it. Otherwise, we assign one label to the event. The order of letters, except for the letter for migration, indicates the order of events. In addition, if selection is associated with selection parameters, we assign a capital letter and, otherwise, a lowercase letter.

### 3.1 Update mechanisms with coupled migration

In this class of update mechanisms, the first event is directly coupled to migration. The first event can be birth or death. If the birth occurs first, one of the patches is selected randomly proportional to the size of the patch (**b**) or randomly proportional to the sum of the selection parameters for birth of that patch (**B**). Next, an individual from the patch is selected purely randomly (**b**) or randomly proportional to its selection parameter for birth (**B**) to produce an identical offspring. The offspring can either stay and substitute one of the individuals in its patch (Fig. 3A) or migrate to one of the neighbouring patches and replace one of the individuals there (Fig. 3B).

If the offspring stays in its own patch, one of the individuals is chosen for death purely at random (**d**) or proportional to its selection parameter for death (**D**) (a common choice is the inverse of the selection parameter for birth). If the offspring migrates to a neighbouring patch, an individual in an adjacent patch is selected in two stages. First, among the neighbouring patches, one patch is selected randomly proportional to the size of the patch (**d**) or randomly proportional to the collective selection parameter for death of the patch (**D**). Finally, one of its individuals is selected for death uniformly at random (**d**) or randomly proportional to its selection parameter for death (**D**).

As an example, consider the update mechanism **MbBDd**. In this update mechanism (i) first, a random patch is selected, (ii) then, from the random patch, one individual is selected proportional to its selection parameter for birth to produce an offspring (iii) next, with a certain probability, the offspring will migrate to one of the adjacent patches or remain in its innate patch. In the former case, one of the neighbouring patches is selected at random as a function of its collective selection parameter for death. (iv) Finally, in the selected patch, which can be either an adjacent patch or the innate patch, one individual dies uniformly at random, and the offspring fills its empty spot.

Based on this procedure, there are 16 different update mechanisms, with birth being the first event; see table 3.1. Similarly, if the first event is death, there are 16 different update mechanisms; see table 3.2. So far, only a few of these mechanisms have been studied in detail. For example, **MBBdd** is adopted in [9, 10, 43]. Not all of the observations made in graphs of individuals with the **Bd** update rule carry over to a graph of subpopulations when the update rule is **MBBdd** [10, 43]. In fact, in the graph of subpopulations, the dynamics and, in particular, the fate of advantageous mutants are highly dependent on the pattern of migration, local population size, and the graph structure itself. Also, applying **MddBB** in the graph of subpopulations reduces the chance of advantageous mutants compared to the equivalent well-mixed population with the update mechanism **dB** [43]. Furthermore, employing **MddBB** in the star of islands in which many subpopulations are connected only via a central subpopulation, it is shown that it is the relative size of the local population in the leaves and the center that determines whether the star of islands is an amplifier, reducer or transient amplifier of selection [13]. In general, in this class of update mechanisms, intuition suggests that the more selection is associated with the selection parameters, the more likely a beneficial mutant will spread through the population.

**Table 3.1:**
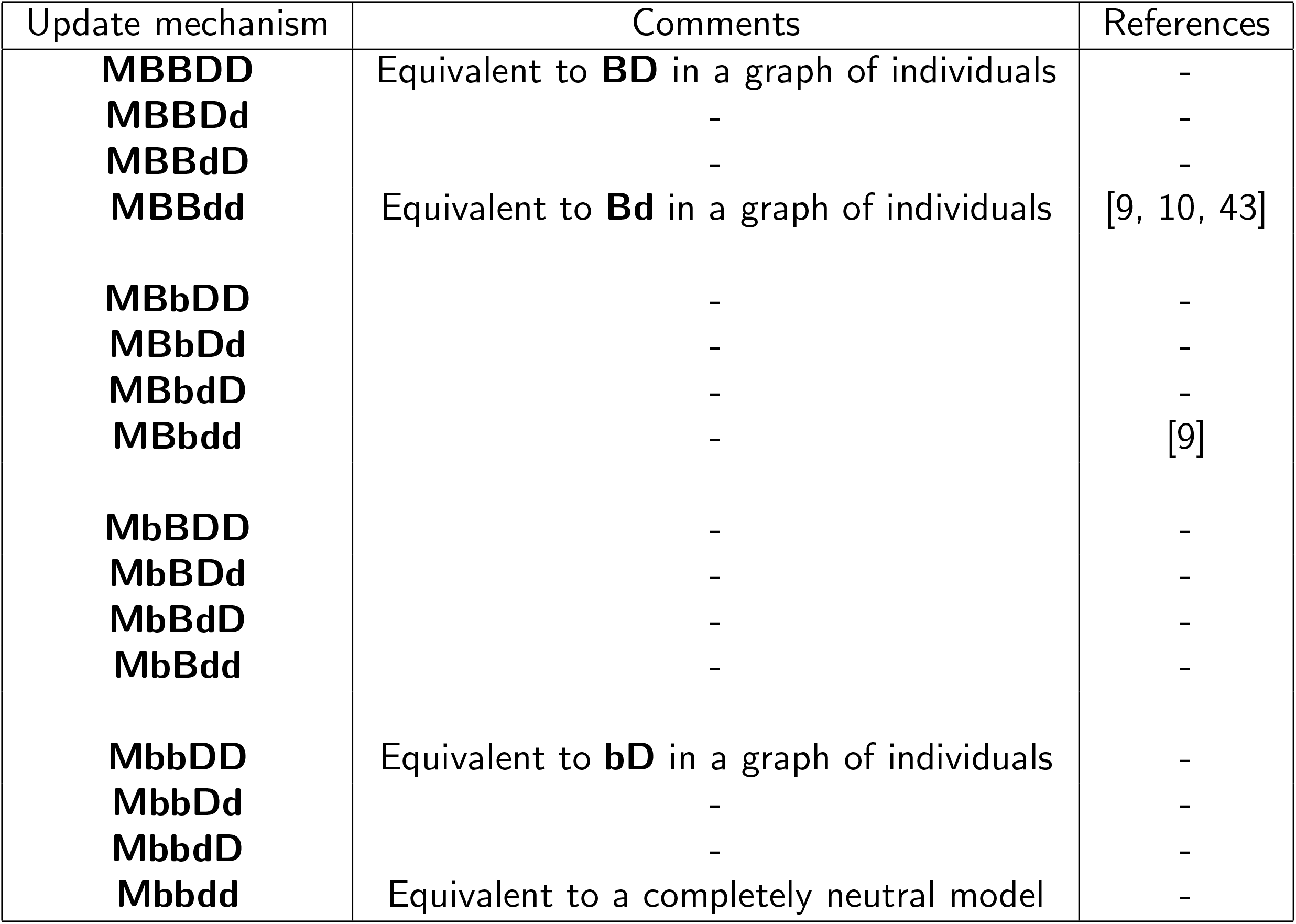
Birth-death processes with migration coupled to reproduction. In all these update mechanisms in graph-structured metapopulations, the individual producing offspring is identified first and the individual to be removed afterwards. In both steps, we can select for the patch and for the individual separately, leading to 16 such update mechanisms.

**Table 3.2:**
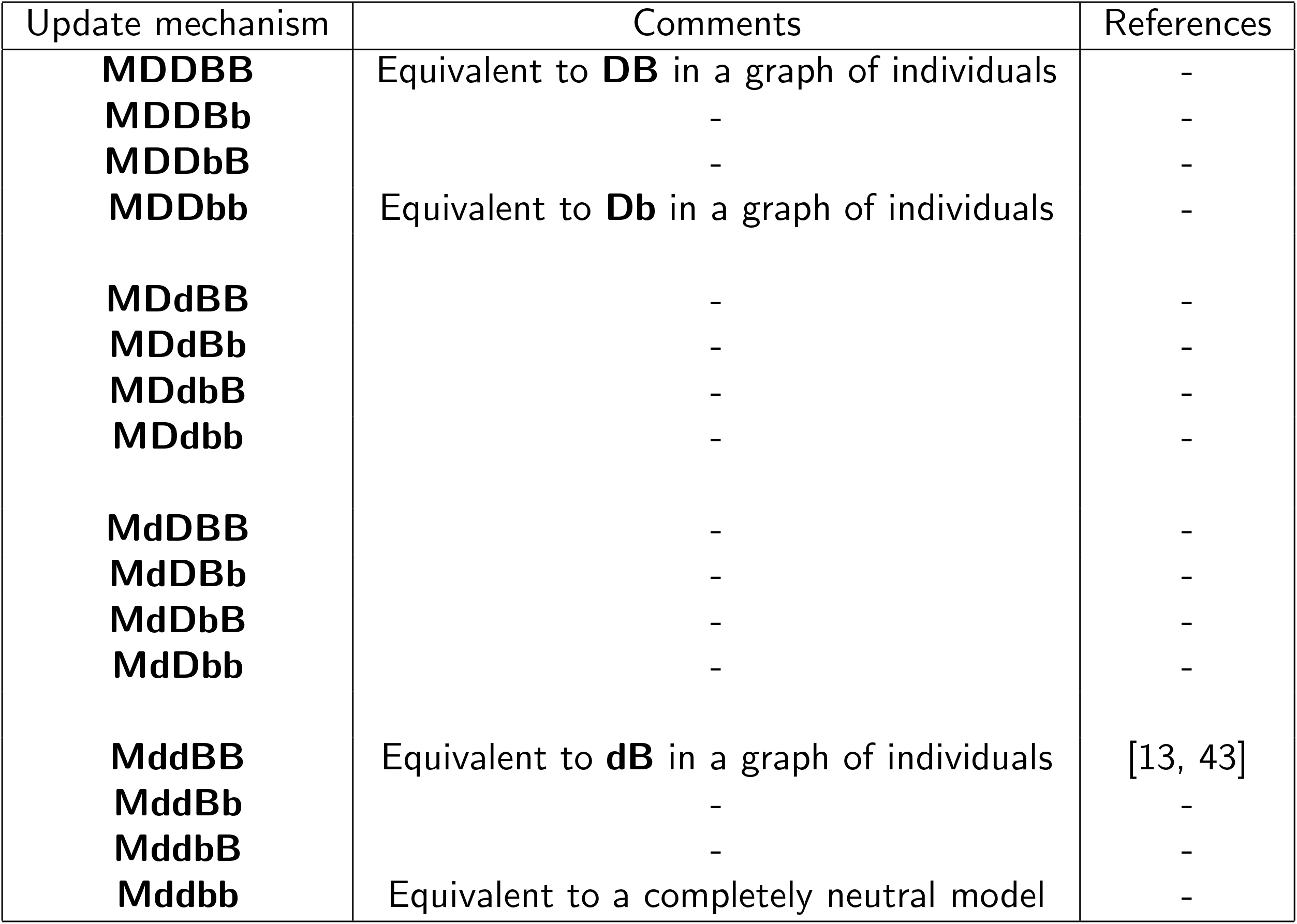
Death-birth processes with migration coupled to death. In all these update mechanisms in graph-structured metapopulations, the individual being removed is identified first and the individual producing offspring afterwards. Again, there are 16 such update mechanisms.

### 3.2 Equivalence to weighted graphs

In the mechanisms with coupled migration, some update mechanisms reduce to simpler ones in a weighted graph of individuals, where the weights of the links that connect individuals in the same subpopulation are different from the weights of the links that connect individuals in different subpopulations (see Fig. 4). For instance, for the update rule **MDDBB**, first, a patch is selected randomly proportional to its collective selection parameter for death. Afterwards, within the patch, an individual is chosen for death randomly proportional to its selection parameter for death. This is equivalent to selecting one individual from the whole population with a probability proportional to its selection parameter for death. Similarly, selection at birth both at the patch and individual levels is equivalent to selecting an individual for birth proportional to its selection parameter for birth from the whole population (for more details go to appendix C).

**Figure 4:**
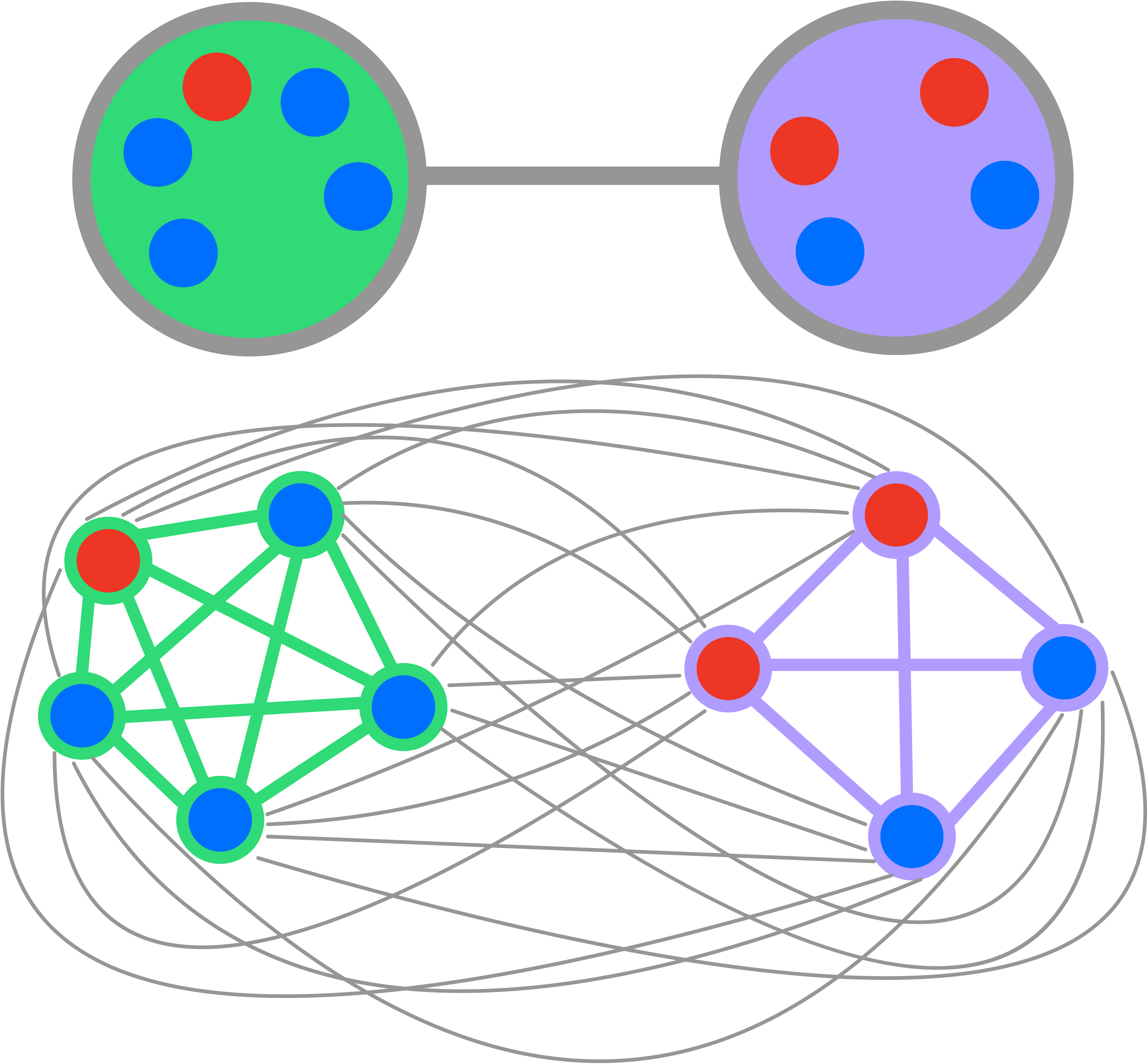
Equivalence of an update mechanism in a graph of subpopulation to an update mechanism in a graph of individuals. In a coupled update mechanism whenever selection on each of the birth and death events both on the patch and individual levels is either purely random or randomly proportional to selection parameters, the update mechanism is equivalent to an update mechanism in an associated weighted graph of individuals. In this weighted graph, the weights of the links that connect local individuals are different from the weights of the links that connect individuals in neighbouring subpopulations and depend on the migration probability as well as local population sizes.

Hence, **MDDBB** on the graph-structured metapopulations can be treated as a **DB** on the equivalent graph of individuals with weighted links between individuals such that the weights of links that connect the individuals belonging to one patch differ from the ones that connect individuals from different patches. In fact, in the coupled update mechanisms whenever selection on each of the birth and death events both on the patch level and individual level is either purely random or randomly proportional to selection parameters (**BB** or **bb**, and **DD** or **dd**), the update mechanism reduces to an update mechanism in an equivalent graph of individuals as mentioned in Tabs. 3.1, 3.2.

### 3.3 Update mechanisms with uncoupled migration

Migration is said to be uncoupled if birth and death events take place within a single patch (Fig. 3A), and migration independently disperses the individuals such that the population size in each patch remains constant (Fig. 3C). In this scenario, we can model migration as follows: with a certain probability, two random individuals from two random connected patches exchange their positions. In this way, the local population size will remain constant.

The first event can be birth or death. If birth happens first, one of the patches is selected randomly (**b**) or proportional to its selection parameter for birth (**B**). From the chosen patch, one individual is selected for birth uniformly at random (**b**) or randomly proportional to its selection parameter for birth (**B**) and produces an identical offspring. Once an individual is selected for birth, one individual is selected for death purely random (**d**) or randomly proportional to a its selection parameter for death (**D**) from the same patch. The new offspring replaces the empty spot of dead individual. In this category, there are eight different update mechanisms. As an example, let us consider **mbBD**. Here, first, a random patch is selected, and in the random patch, an individual is selected randomly proportional to its selection parameter for birth to reproduce. After that, one of the individuals from the same patch is chosen randomly proportional to its selection parameter for death to die. Then, the offspring will fill the empty spot. In this example, there is no patch selection for the second event (death). Hence the second event is indicated by a single letter, **D**.

Similarly, when death happens first, there are eight other update mechanisms. In the update mechanisms **mbbd** and **mddb** where selection is not active in either of the events, the update mechanism is identical to the neutral model i.e. **bd**.

In many popular models of the population genetics literature, migration is assumed to be independent from birth and death [11, 44–51]. However, most of these studies use the Wright-Fisher model as the local update mechanism and are thus not directly equivalent to the models more popular in evolutionary graph theory. If we do not require a strictly constant population size, one can envision many additional update mechanisms, such as the migration of a single individual to another patch. While this appears to be a natural choice in many contexts, it makes the comparison to classical models of Evolutionary Graph Theory more challenging.

## 4 Discussion

Evolutionary Graph Theory is a mathematical framework that has been used to think of the role of population structure in evolutionary dynamics. More recently, empirical scientists have become interested in this framework, but in most of the systems in their focus, the nodes are subpopulations and not individuals. Here, we have classified different classes of update mechanisms on such graph structured metapopulations. We focus on update mechanisms that are natural extensions of the update mechanisms typically used in evolutionary graph theory for graphs of individuals.

Our classification is based on three factors: (i) First, if migration is coupled to reproduction or not. (ii) Second, the order of birth and death events. (iii) Third, how selection acts on the growth and survival of the population. Each of these update mechanisms can result in different dynamics — using different update mechanisms can affect not only the fixation probability and fixation time of newly arising mutations but also other features of the dynamics such as the long term diversity.

The fixation probability in the graphs of individuals under **Bd** when selection for birth is proportional to a selection parameter and **Db** when selection for death is proportional to the inverse of the selection parameter for birth are equivalent in undirected regular graphs [24] and they both follow the isothermal theorem [1]. For more details, see A.1. In addition, in a fully connected graph of individuals, where all individuals are equivalent, and every node in the graph includes a self-loop, meaning that the individual selected for birth can die or the individual selected for death can give birth, **Bd** is equivalent to **dB**, **bD** is equivalent to **Db**, and **BD** is equivalent to **DB**. It is worth-mentioning that in an update mechanism where death is followed by birth, having self-loops in the graphs makes only limited sense if we think of the actual physical death of individuals. However, it is sensible if we think of it in the social setting where death and birth are interpreted as imitating one’s idea or sticking to your own [5]. Furthermore, in a fully-connected graph including self-loops, if selection for death equals the inverse of selection for birth, then the update mechanisms **Bd**, **dB**, **bD**, and **Db** and their corresponding fixation probability is the same as the fixation probability of the well-mixed population under the update mechanism **Bd** [36]. However, in this condition, the fixation probability of an arbitrary graph under **BD** and **DB** is higher than the fixation probability of the corresponding well-mixed population under **Bd**.

In a system where the individuals with a higher selection parameter for birth have a lower selection parameter for death, the more the birth and death are associated with the selection parameters, the fitter individuals have a higher probability to take over the population. This implies that the fixation probability under **BD** is higher than the fixation probability under **Bd** and **bD**. Also the fixation probability under **DB** is higher than the fixation probability under **Db** and **dB**. In addition, the fixation probability of a beneficial mutant in an arbitrary graph under an update mechanism in which selection is global is more than or equal to its fixation probability under an update mechanism in which selection is local [14]. Equality holds for a well-mixed population which includes self-loops meaning that every individual can also replace itself. In a fully connected graph of subpopulations, where the migration rate is symmetric, and the population is homogeneously distributed among the subpopulations, the fixation probability of the update mechanisms such as **MBBdd** that reduce to update mechanisms in a graph of individuals is the same as the fixation probability of the equivalent well-mixed population under update mechanism **Bd** [10].

It is interesting to see how the fixation probability of an arbitrary graph under different update mechanisms changes, as it is known from graphs of individuals that the choice of the update mechanism can be crucial [5]. In the update mechanisms where migration is coupled with death or birth, the fixation probability of an advantageous mutant in an arbitrary graph is higher under some update mechanisms compared to others. As it is shown in Appendix B, if we have two update mechanisms that are exactly the same except that the individual selection for birth or death in one is purely random and in the other is random but associated with selection parameters, the fixation probability of an advantageous mutant under the latter update mechanism is higher than the corresponding fixation probability under the former update mechanism. For example, the update mechanism **MBBDD** does better in fixing the advantageous mutant than **MBbDD**. In addition, one intuitively expects that if selection on the patch level is associated with collective selection parameter, the beneficial mutant has a higher chance of being fixed. Nevertheless, it is not straightforward to prove this. In Appendix B we show this in more detail.

Similarly, as it is shown in Appendix B, for the update mechanisms with uncoupled migration, if we have two update mechanisms that are only different in individual selection on birth or death, the fixation probability of an advantageous mutant under the update mechanism in which individual selection is purely random is smaller than the corresponding fixation probability under the update mechanism in which individual selection is associated with selection parameters. For instance, the fixation probability of an advantageous mutant under **mBBD** is higher than the corresponding fixation probability under **mBBd** in an arbitrary graph.

In the low migration regime, the dynamics of the system under update mechanisms with coupled migration are approximately the same as the dynamics of the system under update mechanisms with uncoupled migration, if selection of patches for birth and death is not associated with the collective selection parameters. This is because in the low migration regime, in both classes, migration is very slow such that it only takes place when the population in each patch is homogeneous. This way, in both classes, migration links homogeneous populations, and it does not change the local dynamics.

In addition, it is interesting to see under which of the coupled or uncoupled update mechanisms, the beneficial mutant has a higher chance of taking over the whole population. Intuitively, under the coupled update mechanism, migration helps to spread beneficial mutants, whereas under uncoupled migration, exchanges the individual between the patches occur randomly and independent of the selection parameters.

We hope that this paper paves the way for future works on the evolutionary dynamics of graph-structured metapopulations. Here, we only classify possible update mechanisms on metapopulations and do not analyze them in detail. However, the update mechanism is a crucial ingredient of evolutionary graph theory and better understanding how it affects evolutionary dynamics in structured metapopulations will be necessary to move the field forward.

## A The transition probabilities for the graph of individuals

Assume a connected graph of individuals where *w_ij_* is the weight of the link connecting node *i* to *j*. The population consists of two types, wild-type A and mutant B. The variable *s_i_* indicates the status of node *i*, *s_i_* = 0 if it is occupied by a wild-type, and *s_i_* = 1 if it is occupied by a mutant. The selection parameter for the birth of the mutant with respect to the wild-type is *r*, and the selection parameter for the death of the mutant with respect to the wild-type is *t*. In an update mechanism where birth is global and death is local, one individual is selected for birth, and then one of its neighbours is chosen for death. The offspring will replace the empty spot. Based on this model, there are two possible transitions: increasing and decreasing the number of mutants by one. The probability 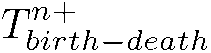 of increasing the number of mutants, 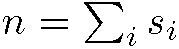 by one is

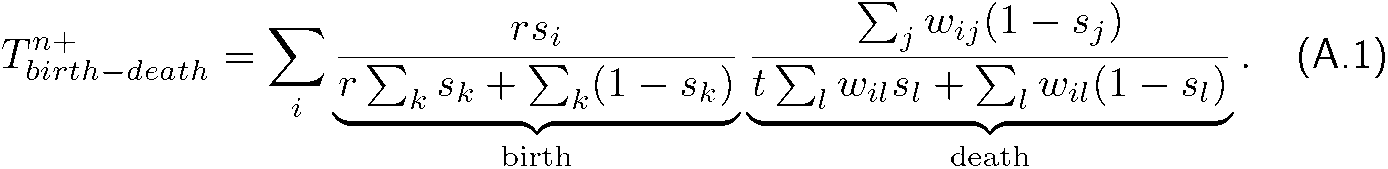

This equation is the summation over all the possibilities that the number of mutants increases. The probability 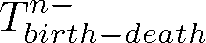 of decreasing the number of mutants *n* by one is

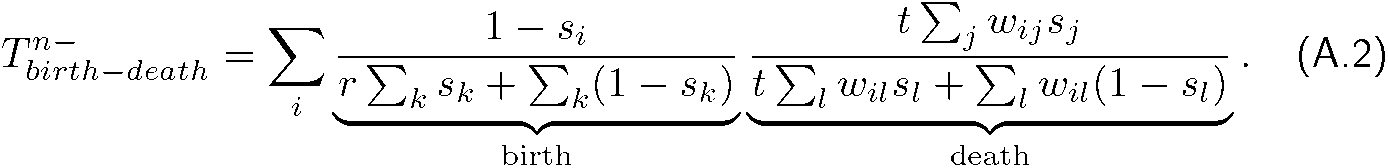

When death is global and birth is local, an individual is selected for death, and then from its neighbour, one individual is selected for birth. In this case the transition probabilities are

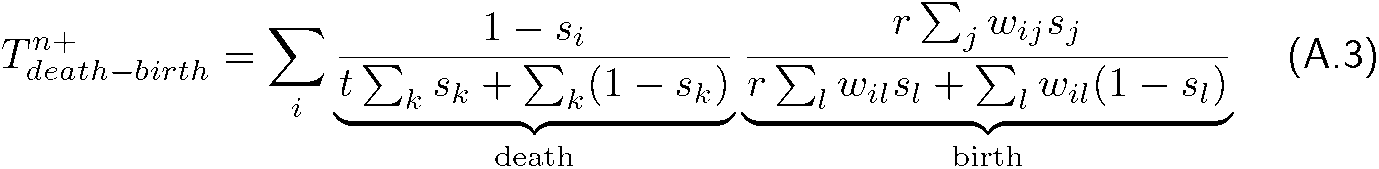

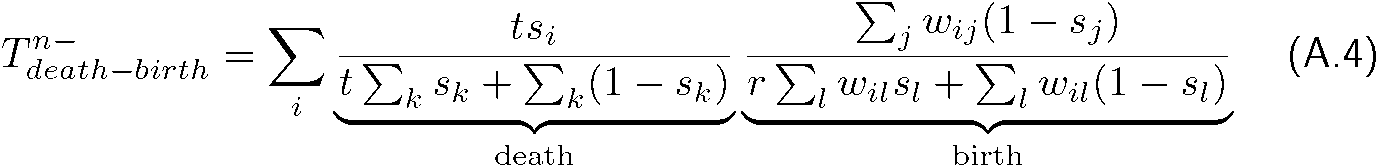

Based on the values of *r* and *t*, the update mechanisms are categorized as follows:

i. In the above equations if *r* ≠ 1 and *t* ≠ 1 there are selection pressures both on the birth and the death. This corresponds to the update mechanism **BD** in which birth is global and death is local, and the update mechanism **DB** in which death is global and death is local.
ii. If *r* ≠ 1 and *t* ≠ 1 birth-death corresponds to update mechanism **Bd** and death-birth corresponds to **dB**.
iii. If *r* ≠ 1 and *t* ≠ 1, birth-death correspond to **bD** and death-birth correspond to **Db**.
iv. If *r* = 1 and *t* = 1 birth-death corresponds to **bd** and death-birth corresponds to **db**.

## A.1 Equivalence of Bd and Db in undirected regular graphs

Using the equations in Appendix A.1, the transition probabilities of an arbitrary graph under update mechanisms **Bd** are

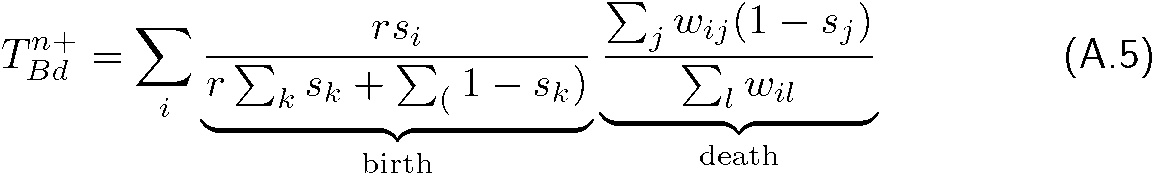

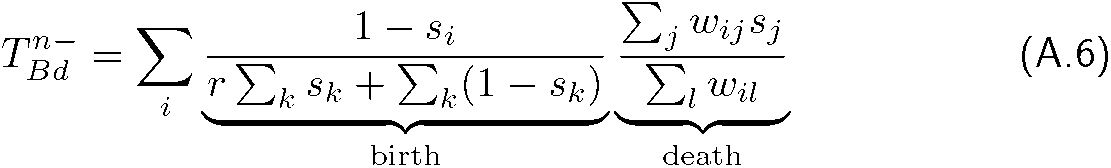

The transition probabilities of an arbitrary graph under update mechanisms **Db** are

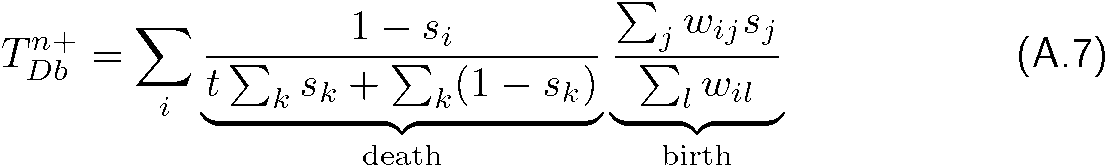

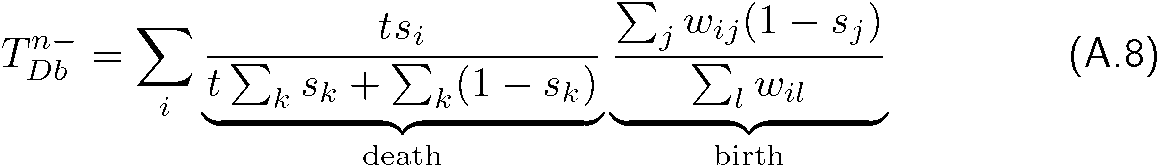

In a regular graph 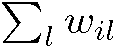 is identical for all the nodes. Thus we can set it as 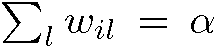. As a result, the transition probabilities for update mechanism **Bd**

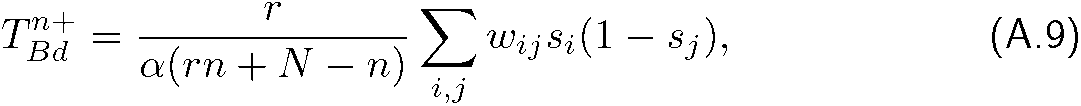

and

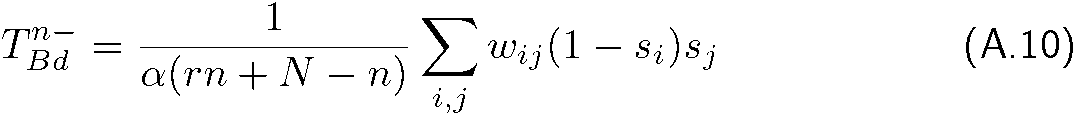

Therefore, 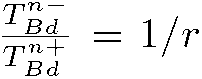 and as a result the fixation probability is 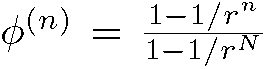. Similarly, the transition probabilities for the update mechanism **Db** simplify

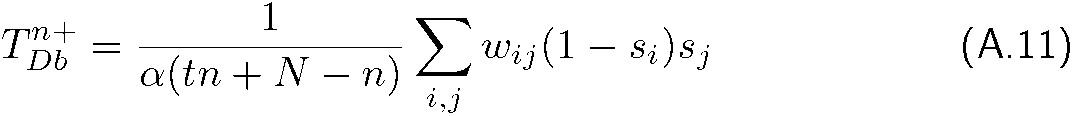

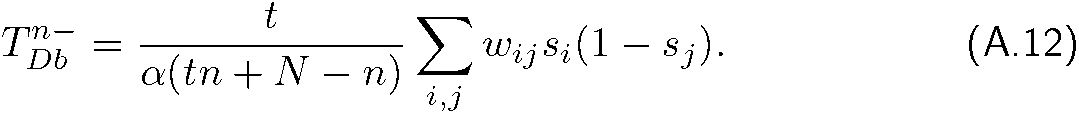

Since 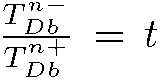, the fixation probability is 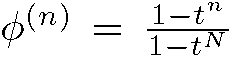. If we set 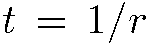 the fixation probability is the same as the fixation probability of the equivalent well-mixed population under the update mechanism Bd.

## B Comparing the fixation probability in structured metapopulations under different update mechanisms

This appendix discusses why some update mechanisms fix beneficial mutants with higher probability. Assume that we have two types of individuals, mutants, and wild-types. The transition probabilities of increasing and decreasing the total number of mutants *n* are given by *T*^*n*+^ and *T*^*n*−^, respectively. Here, we use a mean-field approximation and assume that the transition probabilities are the summation of the transition probabilities for all the possible configurations for a specific number of mutants *n*. The other parameters are described in Tab.B.1. If we start with a single randomly placed mutant in a pure wild-type population, the average fixation probability of a mutant [20] is given by

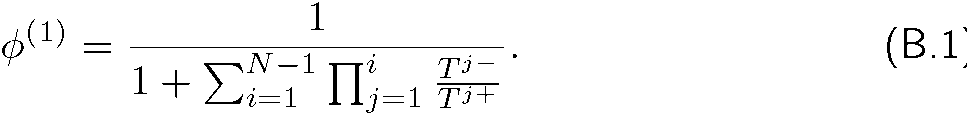

Therefore, in order to investigate how the fixation probabilities in different update rules vary, it is sufficient to compare the transition probabilities. By comparing the transition probabilities, we can see that for both coupled and uncoupled update mechanisms, if two update mechanisms only differ in individual-level selection for either birth or death, the fixation probability of the beneficial mutant under the update mechanism in which individual-selection is associated with selection parameter is higher than the one in which individual-selection is purely random. Here, we compare the transition probabilities of update mechanisms **MBBDD**, **MBbDD**, **MBBDd**. The transition probabilities of increasing and decreasing the total number of mutants, 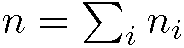 update mechanism **MBBDD** are

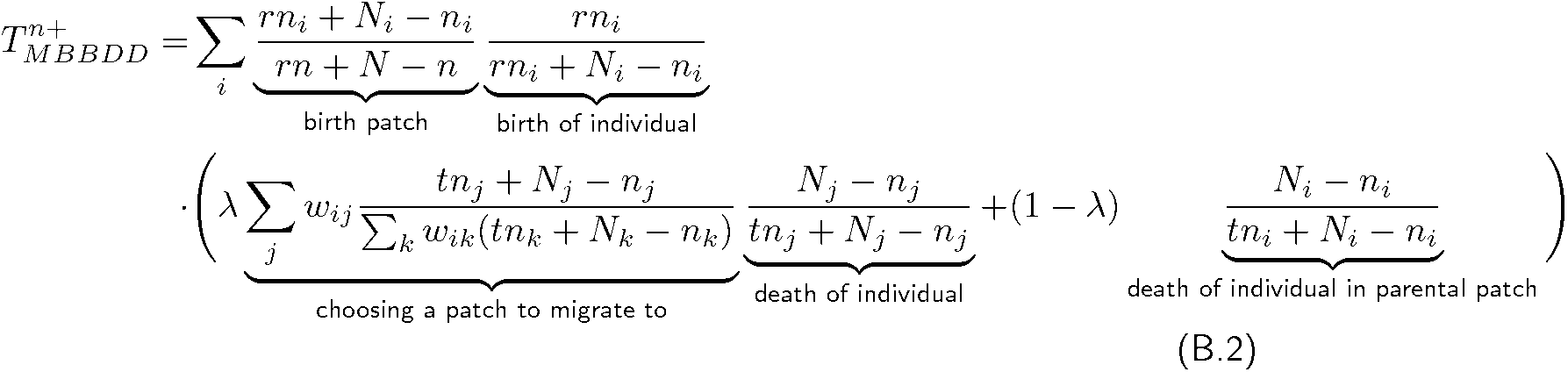

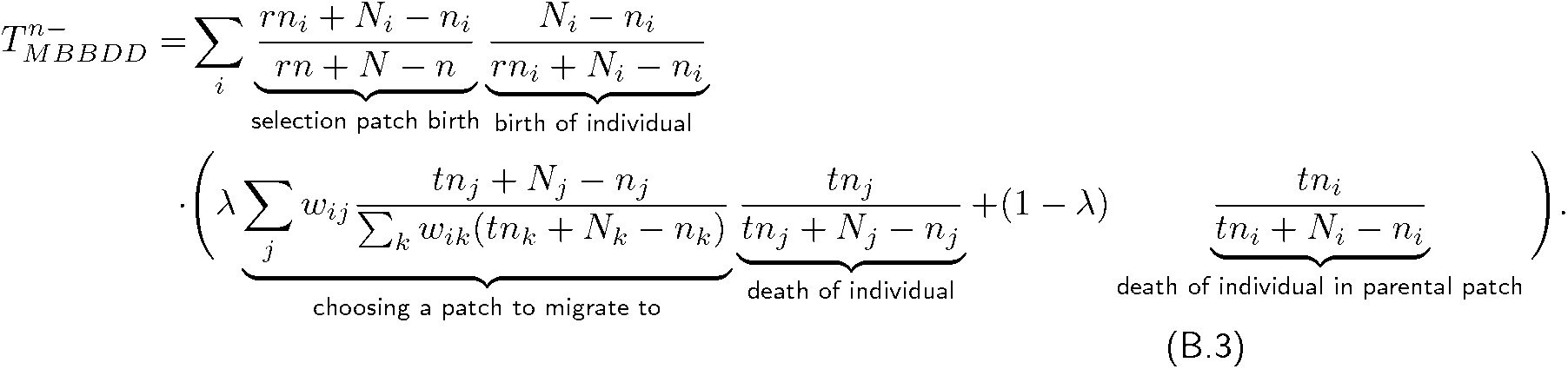

The transition probabilities for the update mechanism **MBbDD** are

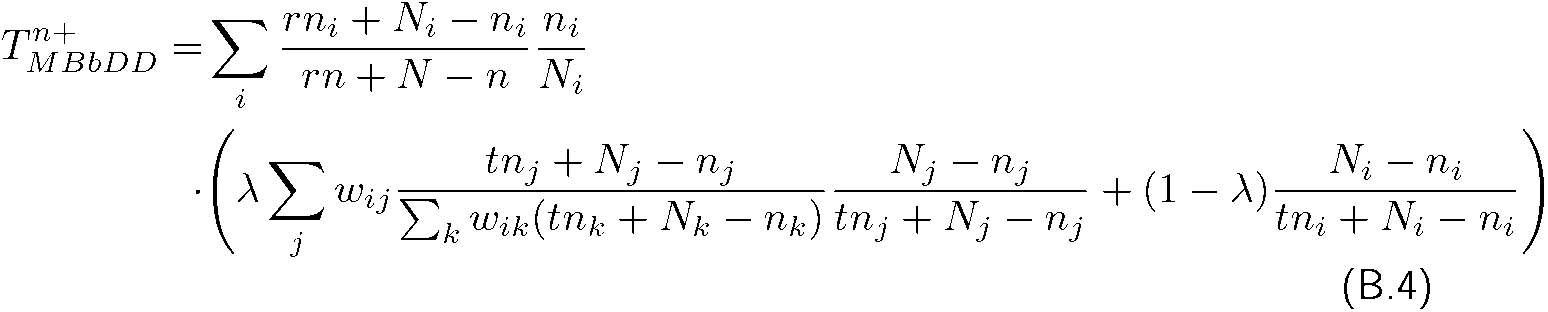

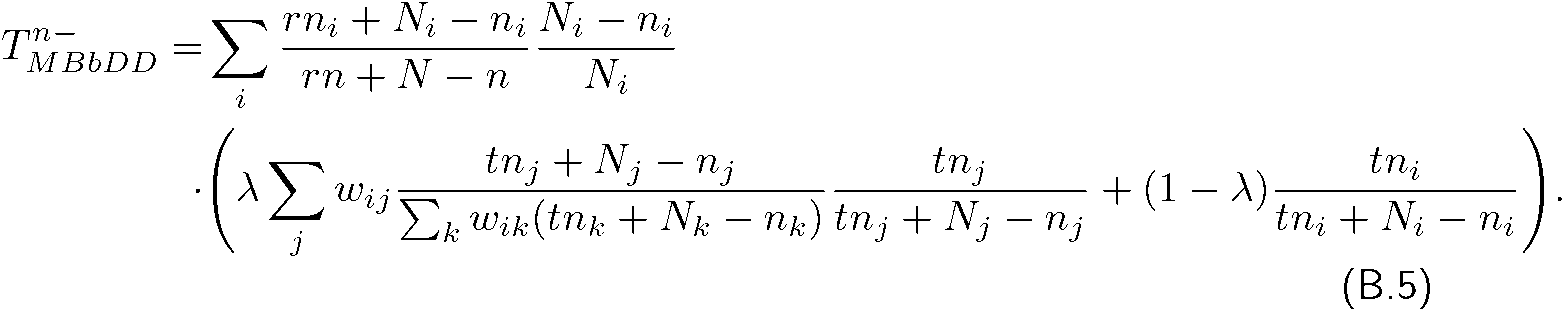

The transition probabilities for the update mechanism **MBBDd** are

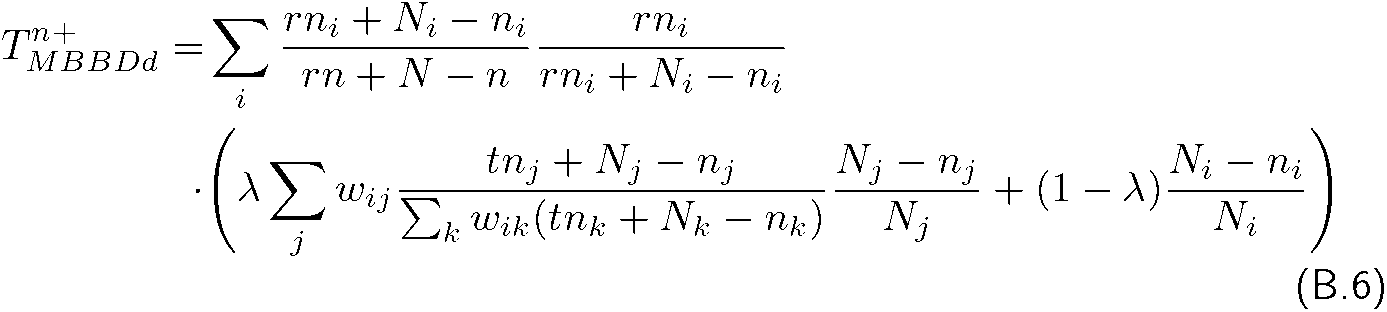

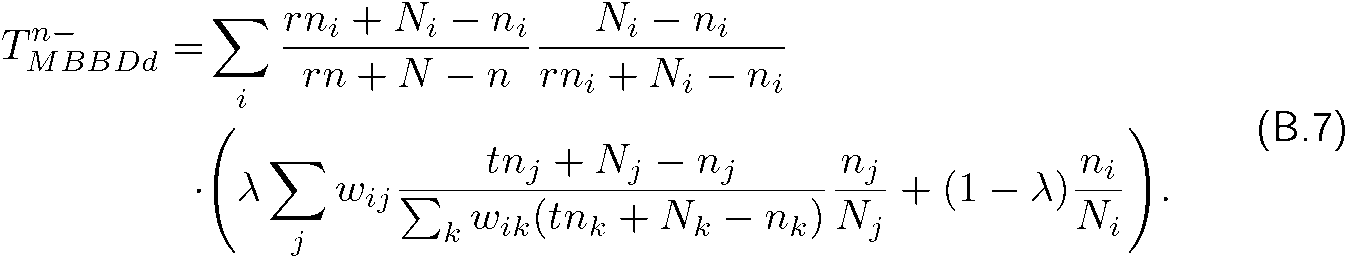

Comparing Eqs. B.2 and B.4, all the terms are the same except the second term which is the probability of choosing a mutant in patch *i* for birth. Since

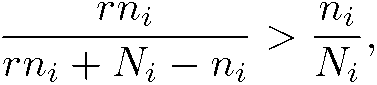

for beneficial mutants, *r* > 1, except for *n_i_* = *N_i_* and *n_i_* = 0 where right hand side and left are equal, that implies

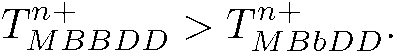

Also if we compare Eqs. B.3 and B.5 since

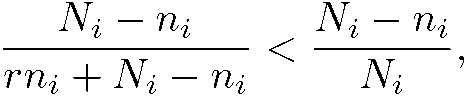

for beneficial mutants, *r* > 1, except for *n_i_* = *N_i_* and *n_i_* = 0 where right hand side and left hand side of the above statement are equal then

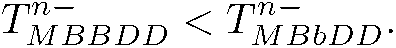

As a result,

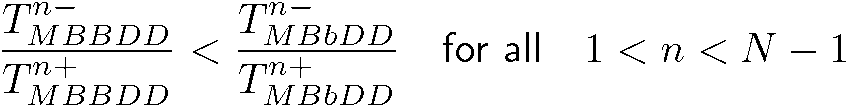

The ratio 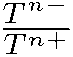 appears in the denominator of Eq. B.1. This implies that the fixation probability of an advantageous mutant under **MBBDD** is higher than the corresponding fixation probability under **MBbDD** for an arbitrary graph, *ϕ_MBBDD_* > *ϕ_MBbDD_*. Similarly, we can show that the fixation probability of a deleterious mutant under **MBBDD** is lower than the corresponding fixation probability under **MBbDD** for an arbitrary graph, *ϕ_MBBDD_* < *ϕ_MBbDD_*.

In addition, comparing Eqs. B.2 and B.6, since

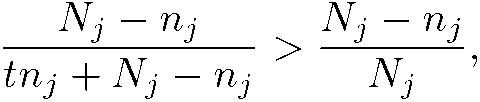

for the beneficial mutants, *t* < 1, we have

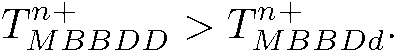

Also by comparing Eqs.B.3 and B.7, we have

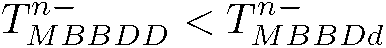

because

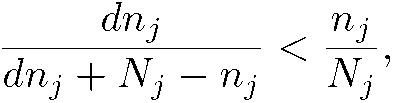

for all 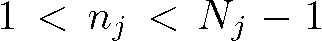 and for 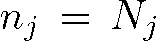 both sides are equal. In conclusion, the fixation probability of an advantageous mutant in an arbitrary graph under **MBBDd** is less than the corresponding fixation probability under **MBBDD**. On the other hand, for deleterious mutants, *ϕ_MBBDd_* > *ϕ_MBBDD_*.

**Table B.1:**
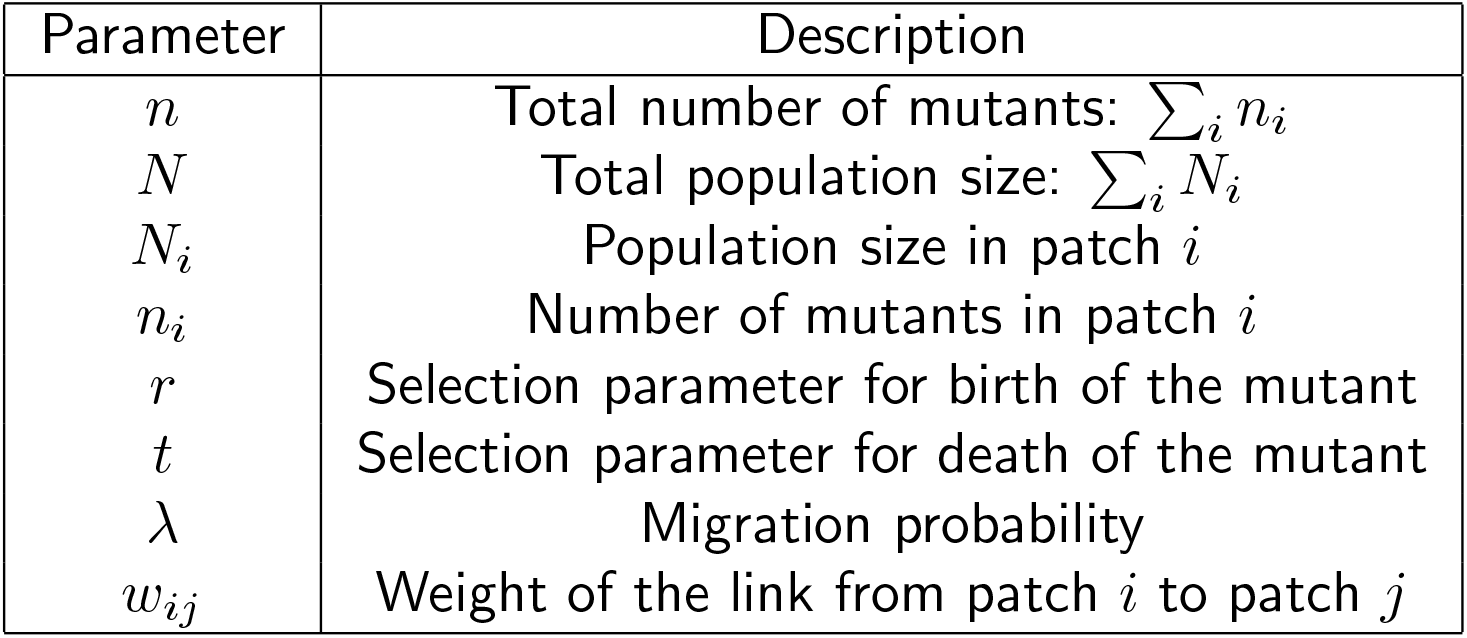
Parameters for graph of subpopulations

Furthermore, we can show in a similar way as above that for beneficial mutants that the fixation probability, *ϕ* of an arbitrary graph under update mechanism **MBBDD**, **MBbDD**, and **MBbDd** has the following relationship with each other,

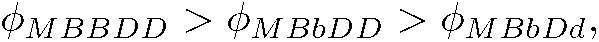

and similarly the relation between the fixation probability of a beneficial mutant for an arbitrary graph under update mechanisms **MBBDD**, **MBBDd**, and **MBbDd** is

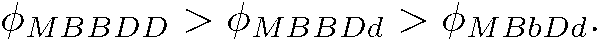

In the above expressions, one cannot simply state that which of the *ϕ_MBBSDd_* and *ϕ_MBbDD_* is higher. The relation between these two values might be dependent on the graph structure.

On the other hand, the relation between the fixation probabilities, *ϕ*, of a deleterious mutant in an arbitrary graph under the update mechanisms, **MBBDD**, **MBbDD**, and **MBbDd** is

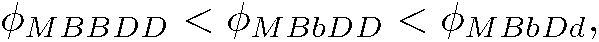

and similarly, we can simply show that the fixation probabilities of a deleterious mutant in an arbitrary graph under the update mechanisms **MBBDD**, **MBBDd**, and **MBbDd** has the following relationship,

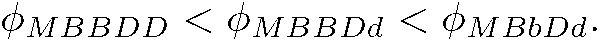

As we see from these equations, for an arbitrary graph, the more an update mechanism is associated with selection parameters, the more the fixation probability of the advantageous mutants and the less the fixation probability of the deleterious mutants.

Comparing the transition probabilities of a beneficial mutant under two update mechanisms that only differ in patch level selection for either birth or death is not straightforward. Intuitively we expect that the update mechanism in which patch level selection is associated with the collective selection parameter of the patch has a higher fixation probability. In the following, we show why one cannot easily infer from the transition probabilities which one is higher. Let us compare the transition probabilities for increasing the population size by one under the update mechanisms **MBBDD** and **MbBDD**.

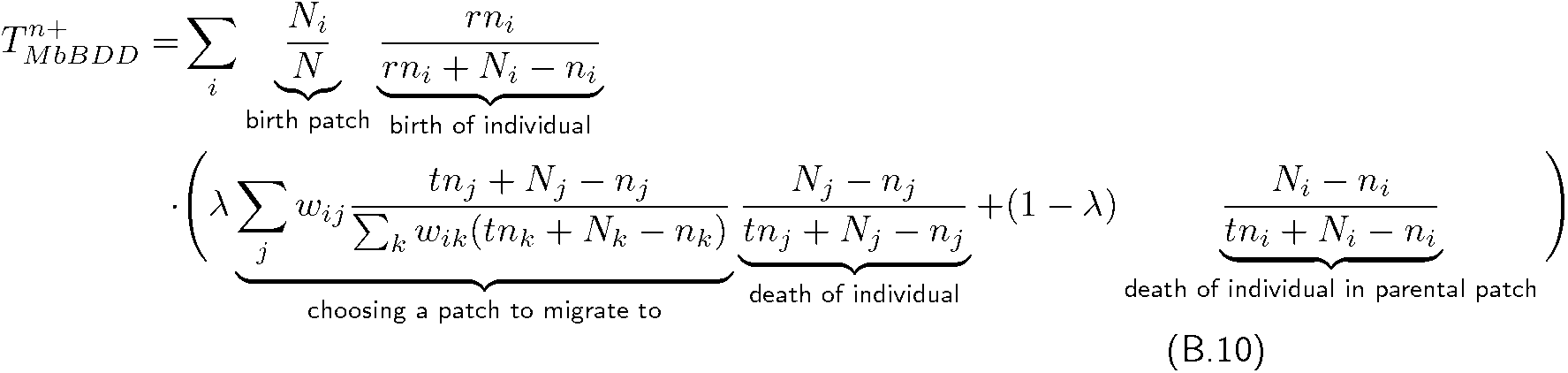

Comparing the equations B.2 and B.10 the only difference is the term for the birth patch. 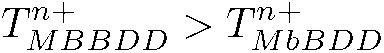 if for all the *i* values

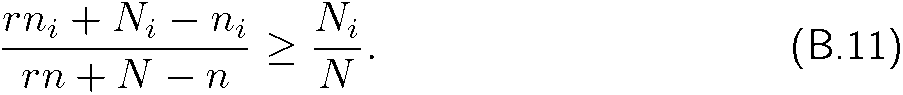

The above relation holds if and only if

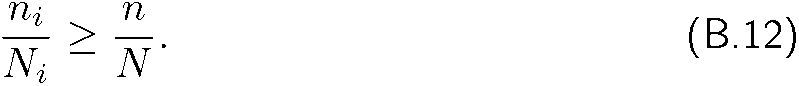

However, the above relation does not always hold, and it depends on the configuration of the population. Hence, it is not easy to compare equations B.2 and B.10.

Similarly, by comparing the transition probabilities for uncoupled update mechanisms, we can see that the fixation probability of an advantageous mutant under update mechanisms in which individual-selection is associated with selection parameters is higher than the corresponding fixation probability under update mechanisms in which individual-selection is independent of selection parameters. The transition probabilities of this update mechanism only include the non-migrative terms because the migrative term does not change the number of mutants in the whole population. The transition probabilities for the update mechanism **mBBD** are

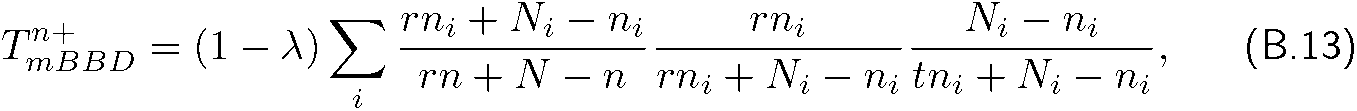

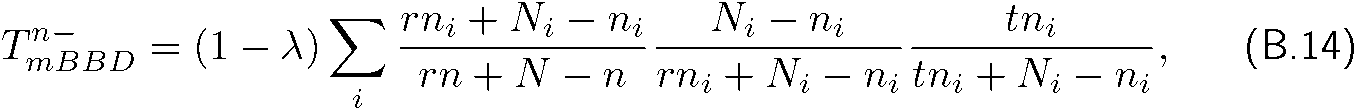

and the transition probabilities for the update mechanism **mBBd** are

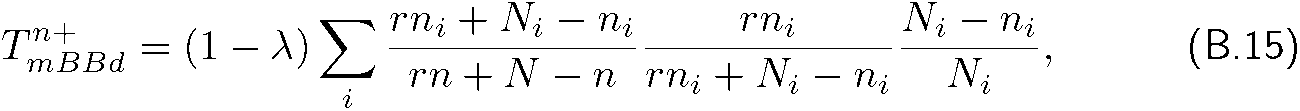

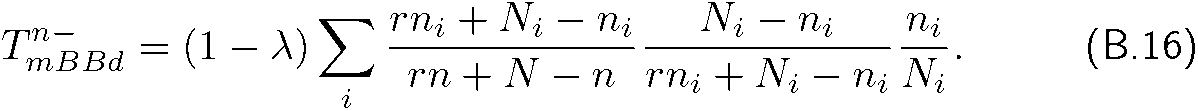

From (Eqs. B.13 and B.15, we can see that for beneficial mutants, *t* < 1,

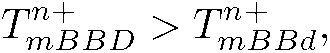

because for 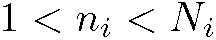 we have

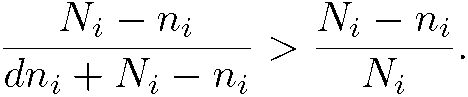

Analogously, for the beneficial mutants we have

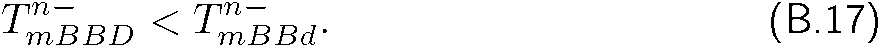

This suggests that the fixation probability of an advantageous mutant under **mBBD** is higher than the corresponding fixation probability under **mBBd**, *ϕ_mBBD_* > *ϕ_mBBd_*. Similarly we can see that for a beneficial mutant

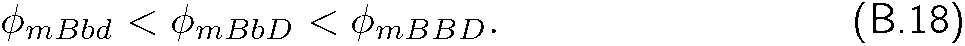

On the other hand, for the deleterious mutants we have an opposite relation between the fixation probabilities:

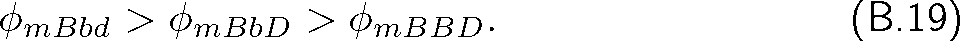

## C Reduction of update mechanisms on graph-structured metapopulations to update mechanisms on graph of individuals

In the mechanisms with coupled migration, some update mechanisms reduce to simpler ones in a weighted graph of individuals, where the weights of the links that connect individuals locally are different from the weights of the links that connect individuals in adjacent subpopulations (see Fig. 4).

In order for an update mechanism to reduce to a simper update mechanism, selection for birth and death should be either associated with selection parameter or not for both patch and individual levels. As an example **MBBdd** reduces to **Bd**. This can be easily shown by transition probabilities; In order to increase the number of mutants by one through selecting one mutant from the patch *i* consists of two parts; first selecting a mutant from patch *i* with probability

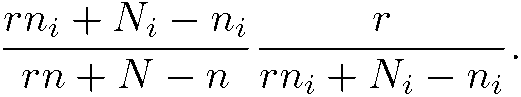

This probability is simplified to 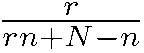 which is equivalent to the probability of selecting one mutant from the whole population regardless of the collective selection parameter of the patches. The second part is choosing one of the wild-type neighbours of the selected mutant for death. The neighbour could be either selected from the parental patch with probability

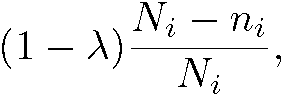

or from the neighbouring patches with probability

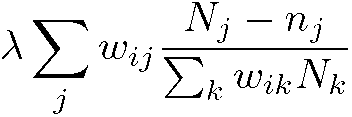

The above two equations for the death probability of a wild-type imply that a graph of patches can be reduced to a graph of individuals. In the equivalent graph of individuals the weight of the link between each two individuals within the patch *i* is 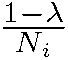 if we take into account self-loops and the weight of the link from an individual from patch *i* to an individual from patch *j* is 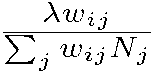. Therefore, the update mechanism **MBBdd** on a graph of sub-populations is equivalent to the update mechanism **Bd** on a graph of individuals in which the weight of the links that connect local individuals differ from the weight of links that connect individuals in different patches. The weight of the links depends on the migration probability as well as the local population sizes.

## References

[1] Lieberman, E., Hauert, C. & Nowak, M. A. Evolutionary dynamics on graphs. Nature 433, 312–316 (2005).

[2] Tkadlec, J., Kaveh, K., Chatterjee, K. & Nowak, M. A. Natural selection of mutants that modify population structure. arXiv preprint arXiv: 2111.10890 (2021).

[3] Gross, T. & Blasius, B. Adaptive coevolutionary networks – a review. Journal of The Royal Society Interface 5, 259–271 (2008).

[4] Perc, M. & Szolnoki, A. Coevolutionary games—a mini review. BioSystems 99, 109–125 (2010).

[5] Hindersin, L. & Traulsen, A. Most undirected random graphs are amplifiers of selection for Birth-death dynamics, but suppressors of selection for death-Birth dynamics. PLoS Computational Biology 11, e1004437 (2015).

[6] Hanski, I. Meta-population dynamics. Nature 396, 41–49 (1998). URL https://www.nature.com/articles/23876. Number: 6706 Publisher: Nature Publishing Group.

[7] Thrall, P. H. et al. Rapid genetic change underpins antagonistic coevolution in a natural host-pathogen metapopulation. Ecology Letters 15, 425–435 (2012).

[8] Eriksson, A., Elias-Wolff, F. & Mehlig, B. Metapopulation dynamics on the brink of extinction. Theoretical Population Biology 83, 101–122 (2013).

[9] Constable, G. W. & McKane, A. J. Population genetics on islands connected by an arbitrary network: An analytic approach. Journal of theoretical biology 358, 149–165 (2014).

[10] Yagoobi, S. & Traulsen, A. Fixation probabilities in network structured metapopulations. Scientific Reports 11, 1–9 (2021).

[11] Marrec, L., Lamberti, I. & Bitbol, A.-F. Toward a universal model for spatially structured populations. Physical Review Letters 127, 218102 (2021).

[12] Zukewich, J., Kurella, V., Doebeli, M. & Hauert, C. Consolidating birth-death and death-birth processes in structured populations. PLoS One 8, e54639 (2013).

[13] Allen, B. et al. Transient amplifiers of selection and reducers of fixation for death-birth updating on graphs. PLoS computational biology 16, e1007529 (2020).

[14] Kaveh, K., Komarova, N. L. & Kohandel, M. Thedualityofspatial death-birth and birth-death processes and limitations of the isothermal theorem. Royal Society Open Science 2 (2015).

[15] Burger, R. Mathematical properties of mutation-selection models. Genetica 102, 279–298 (1998).

[16] Johnson, T. The approach to mutation-selection balance in an infinite asexual population, and the evolution of mutation rates. Proceedings of the Royal Society of London. Series B: Biological Sciences 266, 2389–2397 (1999).

[17] Yagoobi, S., Yousefi, H. & Samani, K. A. Mutation-selection stationary distribution in structured populations. Physical Review E 98, 042301 (2018).

[18] Sharma, N. & Traulsen, A. Suppressors of fixation can increase average fitness beyond amplifiers of selection. Proceedings of the National Academy of Sciences 119, e2205424119 (2022).

[19] Ewens, W. J. Mathematical Population Genetics. I. Theoretical Introduction (Springer, New York, 2004).

[20] Nowak, M. A. Evolutionary dynamics: Exploringtheequationsoflife (Harvard University Press, 2006).

[21] Broom, M. & Rychtar, J. An analysis of the fixation probability of a mutant on special classes of non-directed graphs. Proceedings of the Royal Society A 464, 2609–2627 (2008).

[22] Ohtsuki, H., Hauert, C., Lieberman, E. & Nowak, M. A. A simple rule for the evolution of cooperation on graphs. Nature 441, 502–505 (2006).

[23] Ohtsuki, H. & Nowak, M. A. The replicator equation on graphs. Journal of Theoretical Biology 243, 86–97 (2006).

[24] Antal, T., Redner, S. & Sood, V. Evolutionary dynamics on degree-heterogeneous graphs. Physical Review Letters 96, 188104 (2006).

[25] Sood, V., Antal, T. & Redner, S. Voter models on heterogeneous networks. Physical Review E 77, 13 (2008).

[26] Foo, J., Gunnarsson, E. B., Leder, K. & Sivakoff, D. Dynamicsofadvantageous mutant spread in spatial death-birth and birth-death moran models. arXiv preprint arXiv:2209.11852 (2022).

[27] Kuo, Y. P., Nombela-Arrieta, C. & Carja, O. A theory of evolutionary dynamics on any complex spatial structure. bioRxiv (2021).

[28] Allen, B. et al. Evolutionary dynamics on any population structure. Nature 544, 227–230 (2017).

[29] Tkadlec, J., Pavlogiannis, A., Chatterjee, K. & Nowak, M. A. Limits on amplifiers of natural selection under death-birth updating. PLoS computational biology 16, e1007494 (2020).

[30] Nakamaru, M., Nogami, H. & Iwasa, Y. Score-dependent fertility model for the evolution of cooperation in a lattice. Journal of Theoretical Biology 194, 101–124 (1998).

[31] Nowak, M. A. & May, R. M. Evolutionary games and spatial chaos. Nature 359, 826–829 (1992).

[32] Nakamaru, M., Matsuda, H. & Iwasa, Y. The evolution of cooperation in a lattice-structured population. Journal of Theoretical Biology 184, 65–81 (1997).

[33] Holley, R. A. & Liggett, T. M. Ergodic theorems for weakly interacting infinite systems and the voter model. The annals of probability 643–663 (1975).

[34] Giaimo, S., Arranz, J. & Traulsen, A. Invasion and effective size of graph-structured populations. PLoS Computational Biology 14, e1006559 (2018).

[35] Altrock, P. M. & Traulsen, A. Deterministic evolutionary game dynamics in finite populations. Physical Review E 80, 011909 (2009).

[36] Pattni, K., Broom, M., Rychtář, J. & Silvers, L. J. Evolutionary graph theory revisited: when is an evolutionary process equivalent to the moran process? Proceedings of the Royal Society A: Mathematical, Physical and Engineering Sciences 471, 20150334 (2015).

[37] Masuda, N. Directionality of contact networks suppresses selection pressure in evolutionary dynamics. Journal of Theoretical Biology 258, 323–334 (2009).

[38] Moran, P. A. P. Random processes in genetics. Proceedings of the Cambridge Philosophical Society 54, 60–71 (1958).

[39] Hadjichrysanthou, C., Broom, M. & Rychtář, J. Evolutionary games on star graphs under various updating rules. Dynamic Games and Applications 1, 386–407 (2011).

[40] McAvoy, A. & Allen, B. Fixation probabilities in evolutionary dynamics under weak selection. Journal of Mathematical Biology 82, 1–41 (2021).

[41] Allen, B. & McAvoy, A. A mathematical formalism for natural selection with arbitrary spatial and genetic structure. Journal of mathematical biology 78, 1147–1210 (2019).

[42] Allen, B. & Tarnita, C. E. Measures of success in a class of evolutionary models with fixed population size and structure. Journal of Mathematical Biology (2012).

[43] Houchmandzadeh, B. & Vallade, M. The fixation probability of a beneficial mutation in a geographically structured population. New Journal of Physics 13, 073020 (2011).

[44] Kimura, M. & Weiss, G. The stepping stone model of population structure and the decrease of genetic correlation with distance. Genetics 49, 561–575 (1964).

[45] Wright, S. Evolution in Mendelian populations. Genetics 16, 97–159 (1931).

[46] Maruyama, T. On the fixation probability of mutant genes in a subdivided population. Genetics Research 15, 221–225 (1970).

[47] Maruyama, T. A simple proof that certain quantities are independent of the geographical structure of population. Theoretical Population Biology 5, 148–154 (1974).

[48] Slatkin, M. Fixation probabilities and fixation times in a subdivided population. Evolution 35, 477–488 (1981).

[49] Barton, N. H. The probability of fixation of a favoured allele in a subdivided population. Genetics Research 62, 149–157 (1993).

[50] Whitlock, M. C. Fixation probability and time in subdivided populations. Genetics 164, 767–779 (2003).

[51] Wodarz, D. & Komarova, N. L. Mutant evolution in spatially structured and fragmented expanding populations. Genetics (2020).

